# Reassessing the evolution of strigolactone synthesis and signalling

**DOI:** 10.1101/228320

**Authors:** Catriona Walker, Tom Bennett

## Abstract

Strigolactones (SLs) are an important class of carotenoid-derived signalling molecule in plants, which function both as exogenous signals in the rhizosphere, and as endogenous plant hormones. In flowering plants, SLs are synthesized by a core pathway of four enzymes, and are perceived by the DWARF14 (D14) receptor, leading to degradation of SMAX1-LIKE7 (SMXL7) target proteins in a manner dependent on the SCF^MAX2^ ubiquitin ligase. The evolutionary history of SLs is poorly understood, and it is not clear whether SL synthesis and signalling are present in all land plant lineages, nor when these traits evolved. We have utilized recently-generated genomic and transcriptomic sequences from across the land plant clade to resolve the origin of each known component of SL synthesis and signalling. We show that all enzymes in the core SL synthesis pathway originated at or before the base of land plants, consistent with the previously observed distribution of SLs themselves in land plant lineages. We also show that the late-acting enzyme LATERAL BRANCHING OXIDOREDUCTASE (LBO) is considerably more ancient than previously thought. We perform a detailed phylogenetic analysis of SMXL proteins, and show that specific SL target proteins only arose in flowering plants. We also assess diversity and protein structure in the SMXL family, identifying several previously unknown clades. Overall, our results suggest that SL synthesis is much more ancient than canonical SL signalling, consistent with the idea that SLs first evolved as rhizosphere signals, and were only recruited much later as hormonal signals.

## INTRODUCTION

The ability to tailor growth and development to prevailing environmental conditions is a key feature of plant biology, and has been instrumental in the successful colonization of all terrestrial biospheres by plants. Plant growth is coordinated in space and time through the production and systemic transport of plant hormones; external modulation of these signals allows coupling of environment and development. Strigolactones (SLs) are a class of carotenoid-derived signalling molecules which function endogenously as hormones, while also acting in an exogenous manner as signals in the rhizosphere (reviewed in Waters et al, 2017). SLs play a key role in multiple developmental pathways, including the regulation of shoot branching, lateral root formation and leaf growth. Additionally, the exudation of SLs from the roots into the soil has been shown to be a key factor for the recruitment of arbuscular mycorrhizal (AM) fungi (Akiyama et al, 2005). SLs are particularly associated with soil phosphate levels; SL synthesis is upregulated in low phosphate conditions (Lopez-Raez et al, 2008) and the subsequent recruitment of AM fungi provides the plant with phosphate in exchange for reduced carbon. A proportion of the SLs synthesised within the root are transported into the shoot system, where an inhibitory effect on shoot branching allows the plant to modify shoot system size in direct relation to the availability of soil-borne resources (Kohlen et al, 2011).

In flowering plants (angiosperms), the synthesis of SLs is carried out by a core pathway of four enzymes, which have been characterized in multiple species (reviewed in Waters et al, 2017). The initial substrate all-trans-β-carotene is processed by the carotene isomerase DWARF27 (D27) to 9-cis-β-carotene (Alder et al, 2012), which is subsequently cleaved and modified by two carotenoid cleavage dioxygenases (CCD7 and CCD8) in turn (Alder et al, 2012). The resulting product, carlactone (CL) is active as a SL, but is usually modified by cytochrome P450 enzymes of the MAX1 family to form carlactonoic acid (CLA) or other derivatives (Abe et al, 2014; Zhang et al, 2014). These intermediates are thought to be further processed by an array of enzymes that result in a diverse set of active SL structures (e.g. Brewer et al, 2016). In Arabidopsis, LATERAL BRANCHING OXIDOREDUCTASE (LBO) has been identified as late-acting enzyme that converts CLA to Methyl-CLA (MeCLA), but it assumed further enzymes must also exist, as MeCLA is not an abundant naturally occurring SL in Arabidopsis (Brewer et al, 2016). SL signalling is mediated by the DWARF14 (D14) α/β hydrolase receptor, which binds and enzymatically cleaves SLs, in the process becoming covalently bound to one of the reaction products (Yao et al, 2016; de Saint Germain et al, 2016). This triggers a conformational change in D14 that mediates its interaction with MAX2, an F-box protein that forms part of an SCF ubiquitin ligase complex, which targets proteins for proteolytic degradation (Yao et al, 2016; de Saint Germain et al, 2016). The target proteins of D14 are members of the chaperonin-like SMAX1-LIKE family, specifically the SMAX1-LIKE7/DWARF53 (SMXL7/D53) sub-family. Recruitment of SMXL7 proteins to the signalling complex by active D14 results in the ubiquitination and subsequent degradation of both the D14 and SMXL proteins (Jiang et al, 2013; Zhou et al, 2013; Soundappan et al, 2015). Turnover of SMXL7 proteins allows downstream SL responses to occur, which seem to include both removal of the PIN1 auxin efflux carrier from the plasma membrane of cells in the stem (Soundappan et al, 2015; Liang et al, 2016; Bennett et al, 2016) and increased transcription of BRC1 (Dun et al, 2012; Soundappan et al, 2015; Wang et al, 2015; Seale et al, 2017). SMXL proteins are not DNA-binding transcription factors, but have a well-conserved ERF-Associated Repressive (EAR) motif, and have thus been proposed to act as intermediates in the assembly of repressive transcriptional complexes, via recruitment of TOPLESS family chromatin remodelling complexes (Smith & Li, 2013). There is some evidence for this, particularly in rice (Song et al, 2017), but generally transcriptional responses to SL are very limited (Mashiguchi et al, 2009), and the EAR motif is not absolutely required for SMXL7 function (Liang et al, 2016). The function of SMXL proteins thus remains rather enigmatic, and it possible that they have multiple cellular functions, both transcriptional and non-transcriptional.

The evolution of SL synthesis and signalling has generated equal amounts of interest and confusion. It is clear that this evolution history is not simple, with different components appearing at different points in the evolutionary record (Waters et al, 2017). For instance, with regard to SL synthesis, it has been proposed that D27 arose in the algal ancestors of land plants, CCD7 at the base of the land plant group, CCD8 after the divergence of liverworts and other land plants, MAX1 within the vascular plant group and LBO specifically within seed plants (Delaux et al, 2012; Challis et al, 2013; Brewer et al, 2016; Waters et al, 2017). Outside flowering plants, SL synthesis has been characterized in the moss *Physcomitrella patens*, where CCD7 and CCD8 act consecutively in CL synthesis as in angiosperms (Proust et al, 2011; Decker et al, 2017). There is some uncertainty about which strigolactones are ultimately synthesised by *P. patens*, with recent analysis suggesting only CL is produced, consistent with the lack of MAX1 orthologue in this species (Decker et al, 2017; Yoneyama et al, 2017). In general, conclusions regarding SL synthesis outside the angiosperms are based on very limited sampling of sequences. Conversely, a more exhaustive approach to sampling has recently demonstrated that SL signalling via canonical D14-type SL receptors appears to be a relatively recent innovation within the seed plants (Bythell-Douglas et al, 2017). Understanding the evolution of SL signalling is complicated by the apparent origin of the signalling pathway by duplication of an existing pathway. D14 proteins are closely related to the KARRIKIN-INSENSITIVE2 (KAI2) sub-family of α/β hydrolases, and appear to have arisen by duplication of *KAI2* near the base of land plants followed by gradual neo-functionalization (Bythell-Douglas et al, 2017). KAI2-like proteins are found in charophyte algae, indicating a very ancient origin for KAI2 itself (Bythell-Douglas et al, 2017). In angiosperms, KAI2 perceives smoke-derived karrikin molecules in the environment, but it also assumed to act as a receptor for an as-yet-unidentified endogenous compound (KAI2-Ligand, KL)(reviewed in Waters et al, 2017). Both D14 and KAI2 signalling act through SCF^MAX2^ (Nelson et al, 2011); MAX2 itself has an ancient origin in the algal ancestors of land plants (Bythell-Douglas et al, 2017). SMXL7 proteins are also closely related to the presumptive targets of KAI2 signalling, members of the SUPPRESSOR OF MAX2 1 (SMAX1) sub-family of the SMXL family. There is currently no evidence regarding the evolution of the SMXL family, and the origin of SMAX1 and SMXL7.

The evolution of SLs thus represents something of enigma, but the evidence is currently highly fragmentary, and based on limited sampling from non-representative genomes. In order to try and unravel this mystery, we have exploited recently-generated genomic and transcriptomic sequences from across the land plant clade, to reassess the distribution and evolutionary history of synthesis and signalling components in land plants.

## RESULTS

### Evolution of D27

Lin et al (2009) and Delaux et al (2012) suggested that D27-like proteins are found in both chlorophyte and charophyte algae, implying that D27 proteins should be present in all land plant groups. We identified umambiguous *D27* genes from the fully sequenced genomes of many angiosperms sequences, *Selaginella moellenorfii*, *Physcomitrella patens* and *Marchantia polymorpha*. We also identified sequences from various algal genomes with similarity to *D27*. We obtained sequences similar to *D27* from transcriptomic datasets for all major taxa, with the exception of hornworts. However, in preliminary phylogenetic analyses none of these transcriptome sequences grouped with the set of *D27* sequences from completed genomes. We thus reciprocally BLASTed these new ‘*DWARF27-LIKE1*’ (*D27L1*) sequences against fully sequenced genomes, and from each identified a gene that was more closely related to *D27L1* than *D27*. We also identified more distantly related genes (*D27L2*), which we used as an outgroup. We performed phylogenetic analysis on all these sequences; overall, the resulting topology was not realistic, with unlikely branching orders and poor bootstrap support (Figure 1). The algal sequences did not behave correctly, and grouped with angiosperm *D27L1* sequences, probably due to long-branch attraction. Nevertheless, it is clear that the *D27* and *D27L1* sequences form two coherent land plant clades (Figure 1), which is also evident in the difference between the sequences at their N-terminus (Supplemental data 1). We believe that the most parsimonious explanation for evolution of D27-like proteins is that a single *proto-D27* sequence was present in the algal ancestors of land plants, and that there was a duplication at the base of land plants, leading to the *D27* and *D27L1* lineages. We compared protein identity between D27, D27L1 and algal sequences, across the positions used in our phylogenetic analysis. For every D27 or D27L1 sequence, we calculated *%* identity with each algal protein from non-seed plants, and then averaged across the three algal sequences. We then averaged these averages across either the D27 or D27L1 clade (Supplemental data 2). This analysis suggests that D27L1 better preserves the ancestral protein structure of the algal D27-like proteins (58.2% average identity) and that D27 proteins represent an innovation in protein structure relative to the algal proteins (42.2% average identity). This tentatively suggests that eu-D27 function arose at the base of land plants. Our failure to identify *D27* sequences from transcriptome assemblies suggests *D27* is either expressed at very low levels, or in a spatially restricted manner, such that it is not represented in these assemblies.

**Figure 1:**
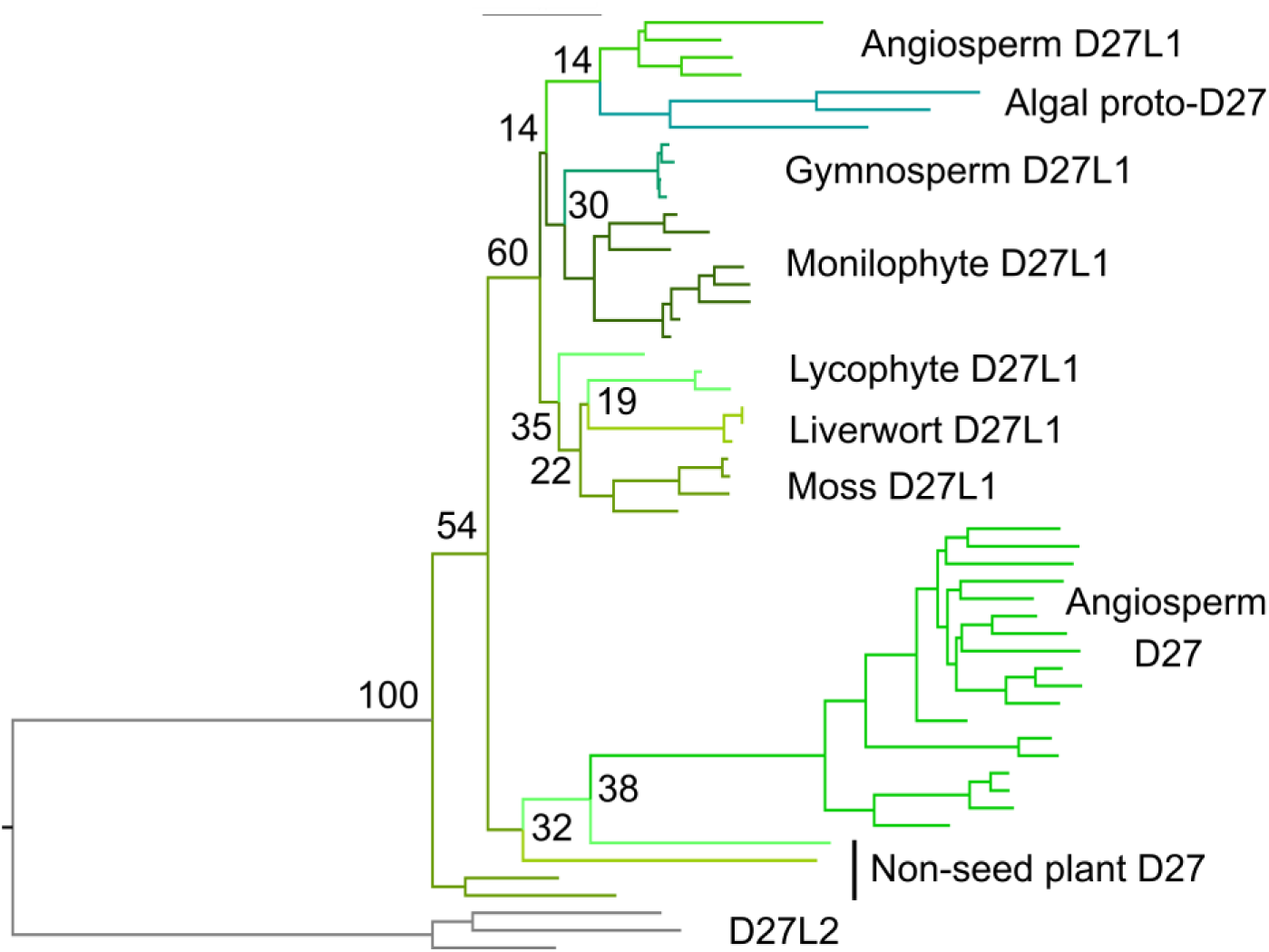
Evolution of D27. Nucleotide-level phylogenetic analysis implemented in PhyML on the *D27* family (54 sequences, 561 characters). The tree was rooted with algal sequences. Phylogram showing the 'most likely’ tree, including bootstrap values at key nodes.

### Evolution of CCD7

We identified *CCD7*-like sequences from chlorophytes, charophytes and the major land plant groups, with the exception of hornworts and monilophytes (Supplemental data 3). Phylogenetic analysis of the broader *CCD* family shows these sequences form a monophyletic group (Supplemental data 4), but as Delaux et al (2012) showed, the algal sequences are quite distinct from land plant sequences, and the proteins may have rather different substrate specificity. The lack of sequences from monilophytes is somewhat surprising, but in general we only identified 6 *CCD7* sequences from transcriptome assemblies, suggesting that *CCD7* expression is generally low or spatially restricted in land plants, and that the lack of sequences from monilophytes is a sampling error. Phylogenetic analysis suggests the evolution history of *CCD7* is relatively simple, with no major duplications present in the family, implying a strong pressure to maintain *CCD7* as a single copy gene (Figure 2).

**Figure 2:**
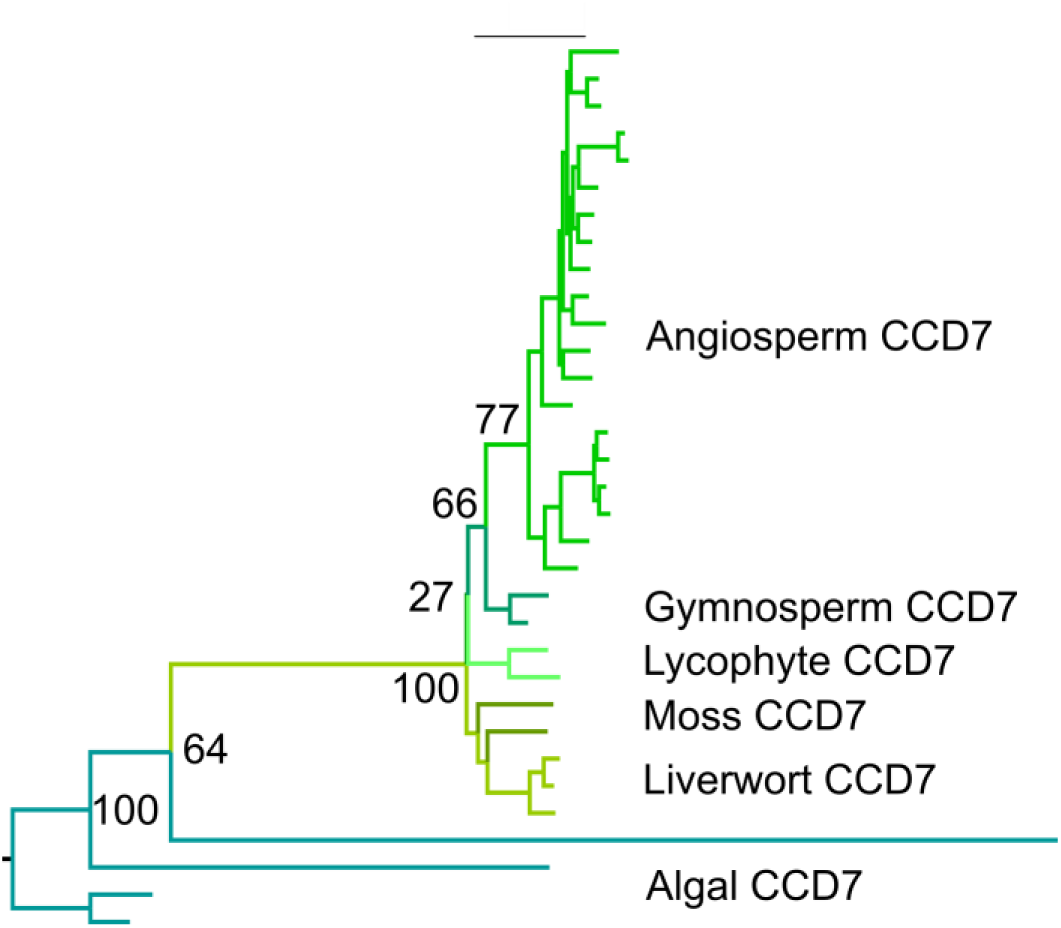
Evolution of CCD7. Nucleotide-level phylogenetic analysis implemented in PhyML on the *CCD7* family (33 sequences, 1599 characters). The tree was rooted with algal sequences. Phylogram showing the ‘most likely’ tree with bootstrap values at key nodes.

### Evolution of CCD8

Previous work has suggested that *CCD8* is absent from liverworts, leaving questions open as to the relative timing of *CCD8* evolution (Waters et al, 2017). However, while *CCD8* is indeed absent from the completed genome of *Marchantia polymorpha*, we obtained multiple unambiguous *CCD8* sequences from other liverworts; *M. polymorpha* is thus an exception, rather than the rule. Indeed, we obtained unambiguous *CCD8* sequences from all the major land plant groups (including hornworts), and CCD8-like sequences from chlorophyte algae (Supplemental data 6); phylogenetic analysis shows that these are all part of a monophyletic clade (Supplemental data 5). As with *CCD7*, phylogenetic analysis suggests evolution of the *CCD8* family is relatively simple; again, there are no major duplications in the family, and *CCD8* is mostly present as single copy gene (Figure 3).

**Figure 3:**
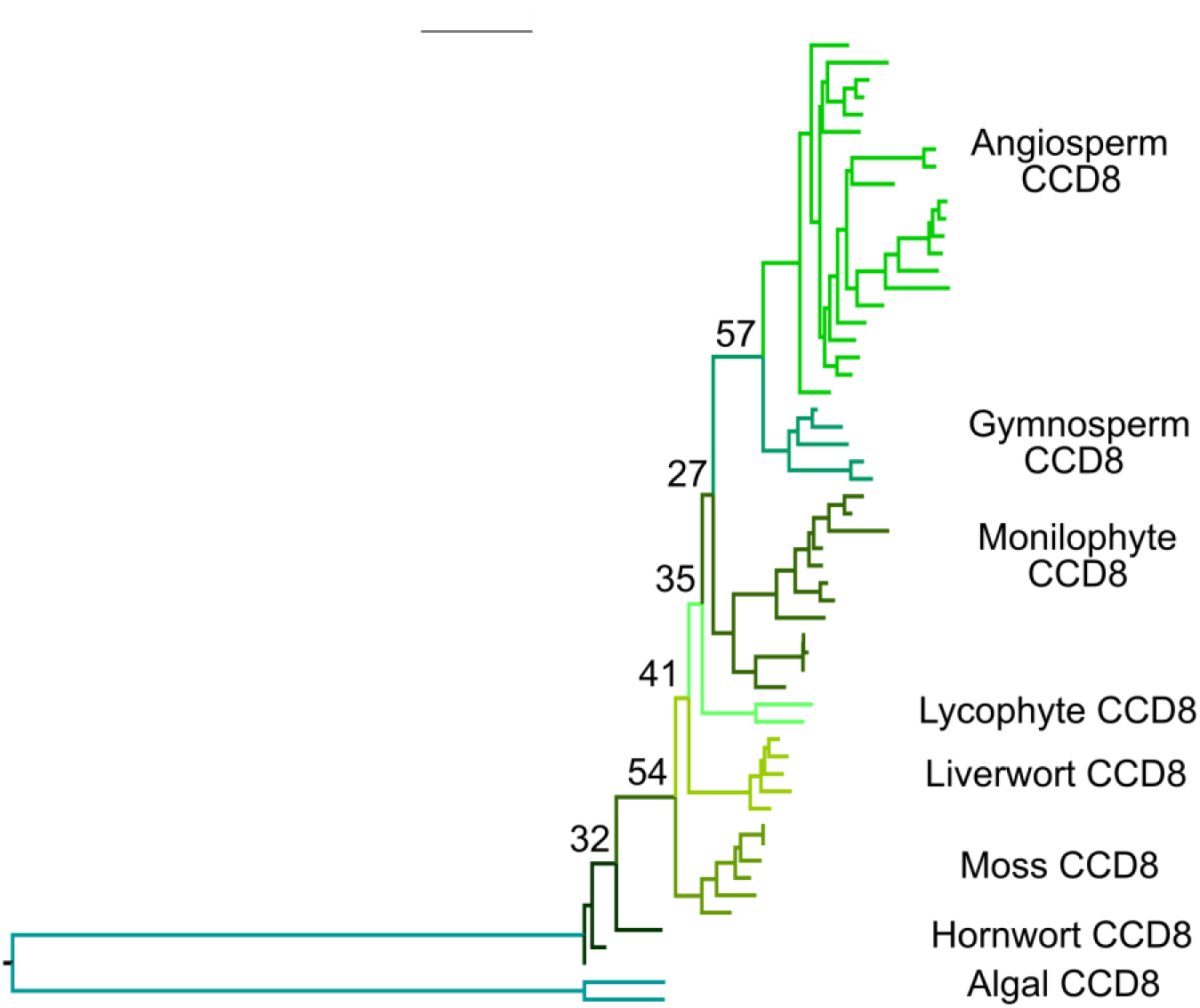
Evolution of CCD8. Nucleotide-level phylogenetic analysis implemented in PhyML on the *CCD8* family (56 sequences, 1449 characters). The tree was rooted with algal sequences. Phylogram showing the ‘most likely’ tree with bootstrap values at key nodes.

### Evolution of MAX1

The evolutionary history of the MAX1 family was previously surveyed by Challis et al (2013), but no *MAX1*-like sequence was identified in *P. patens*, leading to uncertainty regarding the evolutionary origin of the *MAX1*-function in SL synthesis. We identified unambiguous MAX1 sequences from all major land plant groups except hornworts, and a *MAX1*-like sequence from *Klebsormidium nitens*, though we did not obtain any sequences from chlorophyte algae (Supplemental data 7). This suggests a much earlier origin of *MAX1* than previously apparent. While *P. patens* does indeed have no *MAX1*, other mosses possess copies of *MAX1*, as do most liverworts - though as with *CCD8, M. polymorpha* does not (Figure 4). As with the other enzymes, there are no major duplications in the *MAX1* family, which is present as a single copy gene in most species (Figure 4).

**Figure 4:**
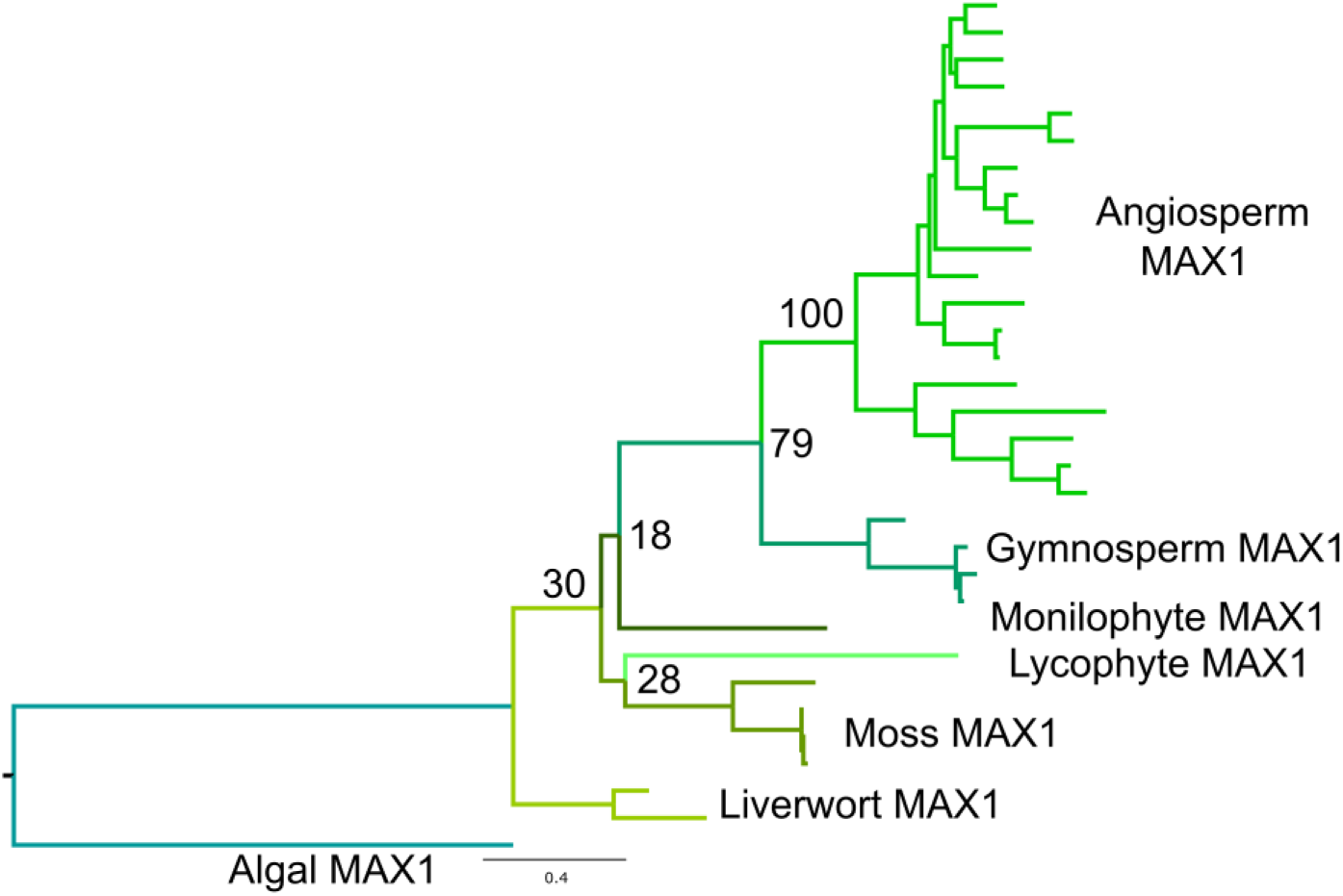
Evolution of MAX1. Nucleotide-level phylogenetic analysis implemented in PhyML on the *MAX1* family (32 sequences, 1425 characters). The tree was rooted with an algal sequence. Phylogram showing the ‘most likely’ tree with bootstrap values at key nodes.

### Evolution of LBO

Based on a relatively simple phylogeny, Brewer et al (2016) concluded that since LBO-like proteins were not present in *Physcomitrella patens* and *Selaginella moellendorfii*, LBO likely represented a seed plant innovation. However, this approach used two non-representative genomes to make conclusions regarding all non-seed plants. We reinvestigated the evolution of *LBO* using a broad sampling approach, and identified a clade of proteins present in all land plants, which contained Arabidopsis LBO. This clade contained the previously described DOXC53 (LBO) and uncharacterized DOXC54 angiosperm clades, and equivalent sequences from gymnosperms (Brewer et al, 2016). We named the DOXC54 clade RELATED TO STRIGOLACTONE SYNTHESIS (RSS). These two clades appear to have arisen from an ancestral gene present in single copy in non-seed plants, by a duplication at the base of seed plants (Figure 5; Supplemental data 8). Brewer et al (2016) suggested that LBO activity arose in seed plants, which would imply the *LBO* lineage has neo-functionalized after the duplication in seed plants. However, the shorter branch lengths in *LBO* relative to *RSS* suggested that LBO was more likely to retain the original function of the proto-LBO proteins than RSS. We thus compared protein sequence between angiosperm/gymnosperm LBO and RSS proteins and the non-seed plant proto-LBO proteins, across the positions used in our phylogenetic analysis. For every seed plant LBO/RSS sequence, we calculated % identity with each protein from non-seed plants. For each seed plant sequence, we then calculated the average % identity with non-seed plant proto-LBO proteins. Finally, we then averaged these averages across either the LBO or RSS clade (Supplemental data 9). On average LBO proteins were 47.1% identical to non-seed plant proto-LBO proteins, while RSS proteins were only 45.6% identical. This tentatively suggests that LBO proteins are more likely to retain the original function of proto-LBO proteins than RSS proteins.

**Figure 5:**
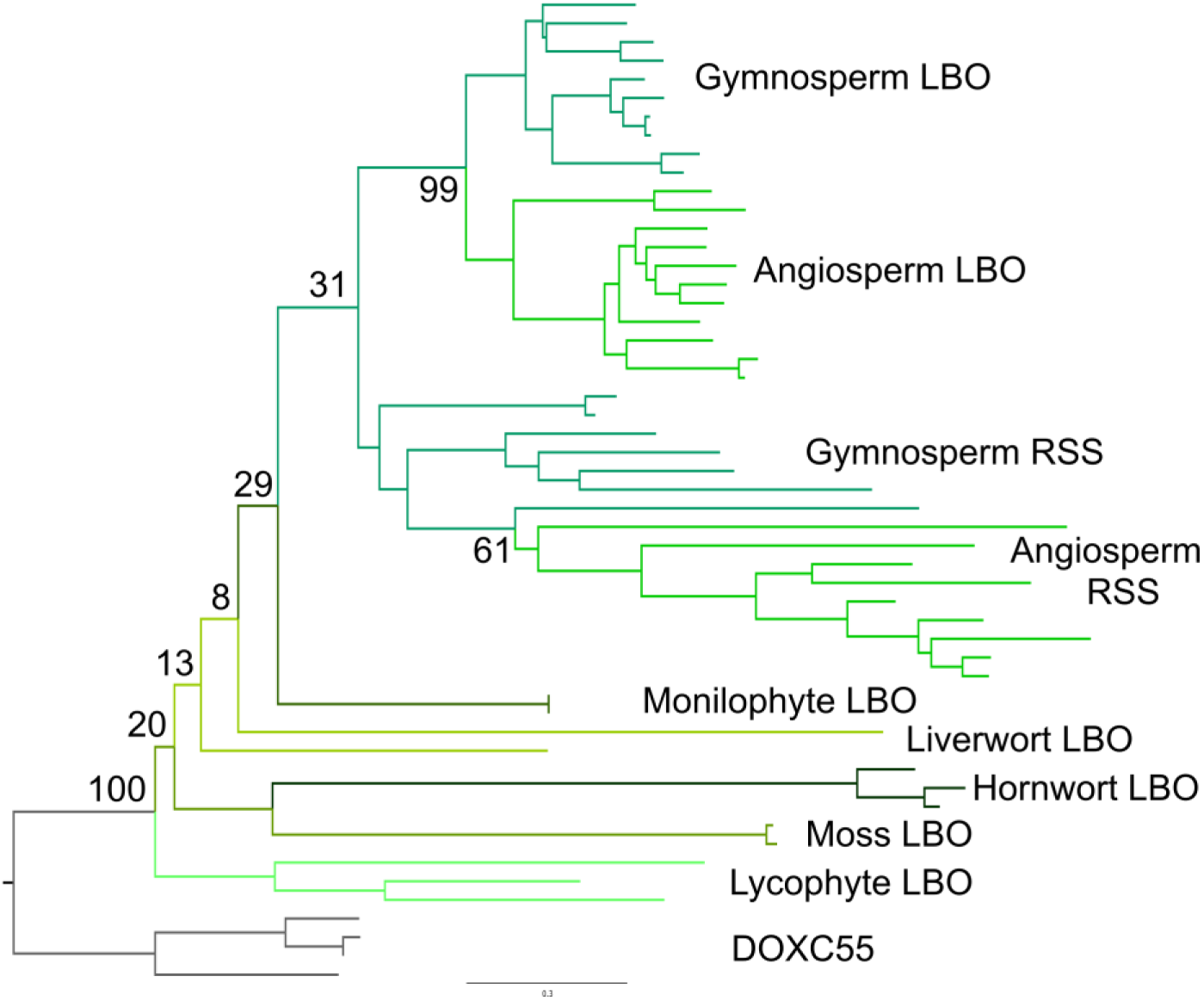
Evolution of LBO. Nucleotide-level phylogenetic analysis implemented in PhyML on the *LBO* family (53 sequences, 852 characters). The tree was rooted with sequences from the DOXC55 clade. Phylogram showing the ‘most likely’ tree with bootstrap values at key nodes.

### SMXL proteins are present throughout land plants but not in algae

We next turned our attention to understanding the evolution of SL signalling. We have recently examined the evolution of SL receptors in the *D14/KAI2* family, and shown that SL perception probably evolved gradually by neo-functionalization of KAI2-like receptors (Bythell-Douglas et al, 2017). We also showed that MAX2 is a deeply conserved protein in charophyte algae and land plants (Bythell-Douglas et al, 2017). However, very little is known regarding the evolution of the SMXL family proteins that are the proteolytic targets of SL and KL signalling. In order to understand *SMXL* evolution, we obtained 226 *SMXL* sequences from 99 species (Supplemental data 10). We identified unambiguous *SMXL* sequences in all major land plant groups, but not in any of the chlorophyte or charophyte genome/transcriptome dataset (Table 1). Preliminary phylogenetic analyses indicated a more complex evolutionary history than the SL synthesis enzymes, and placed *SMXL* family members into clear taxon-specific clades (Table 1). We identified 1 *SMXL* clade in each of the liverworts (*SMXLA*), mosses, hornworts (*SMXLD*), lycophytes and monilophytes (*SMXLJ*) (Table 1). In mosses of the Bryopsida, there are two distinct *SMXL* sub-clades (*SMXLB* and *SMXLC),* but only a single *SMXL* clade (resembling *SMXLB*) is present in the early-diverging Sphagnopsida lineage. These results are consistent with the evolution of the *D14/KAI2* family in mosses, where four *D14/KAI2* sub-clades are present in the Bryopsida, but only two in the Sphagnopsida (Bythell-Douglas et al, 2017). Collectively, these data suggest that a whole genome duplication may have occurred at the base of the Bryopsidan lineage. Similarly, in the lycophyte group, there are two *SMXL* sub-clades present in the Selaginellales (*SMXLG* and *SMXLH*), but only one in the Lycopodiales (*SMXLE*) and Isoetales (*SMXLF*). Although they likely form a monophyletic clade, there is little resemblance between SMXLE, SMXLF, SMXLG or SMXLH. This degree of sequence divergence within the lycophytes is consistent with our previous observations of D14/KAI2 and PIN protein family members (Bythell-Douglas et al, 2017; Bennett et al, 2014). We detected two distinct clades of *SMXL* proteins in gymnosperms and at least four distinct clades in angiosperms (Table 1).

**Table 1:**
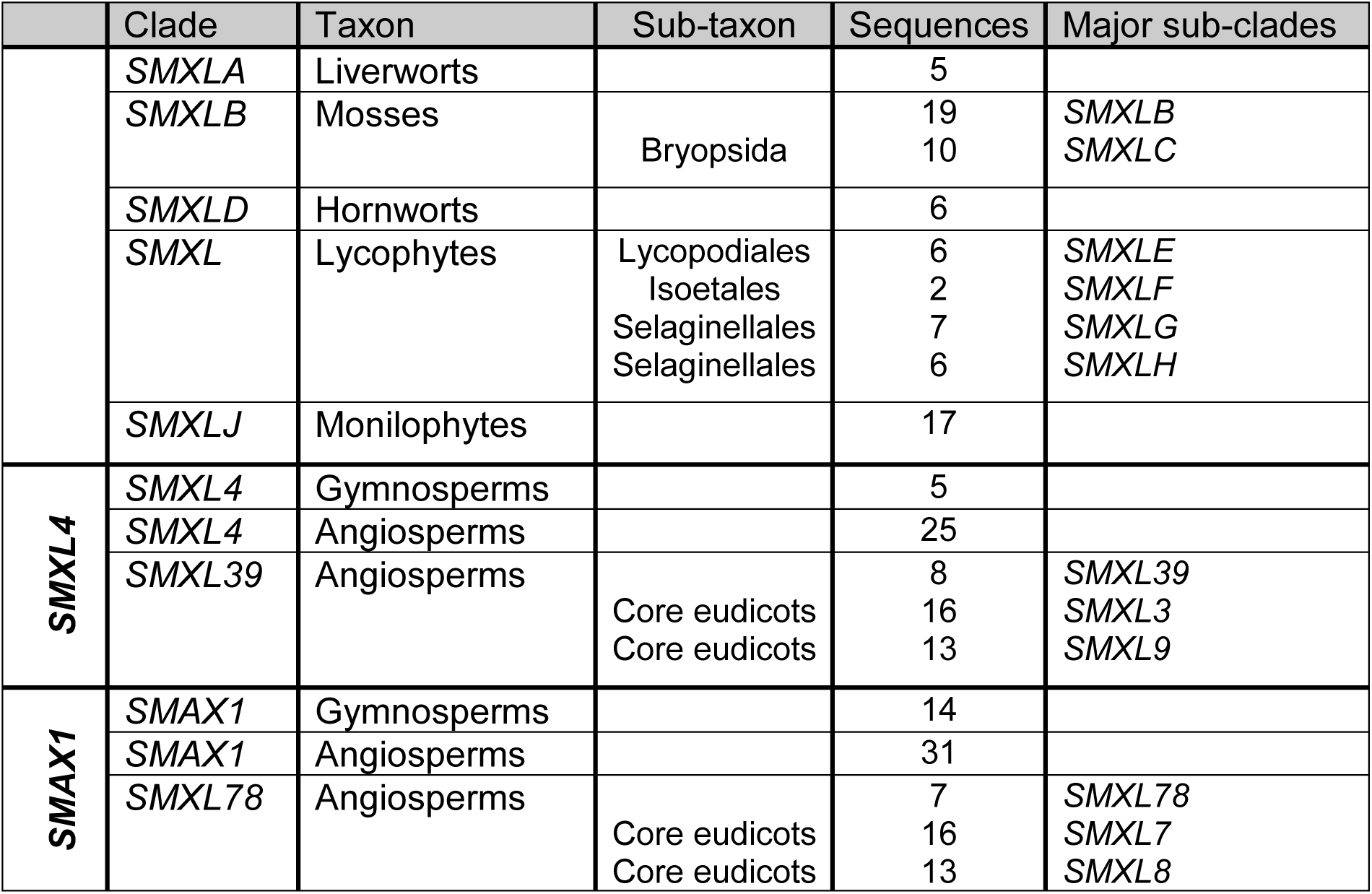
**Major clades in the *SMXL* family** Table showing major clades in the *SMXL* family, as defined at the level of major taxonomic groups. Almost all sequences in the family unambiguously group into one of these clades. Within some clades there are major sub-clades where the lineage has been duplicated; these are listed at the right. Our analysis suggests that seed plant *SMXL* proteins group into two super-clades, *SMAX1* and *SMXL4*, as indicated on the left of the table.

### Diversification of SMXL proteins in the seed plant lineage

To understand the interrelationship of these clades, we reconstructed the evolution of the family using maximum likelihood approaches. All analyses confirm the composition of the major clades in Table 1, and suggest a basic topology for the *SMXL* family. In the absence of an obvious algal outgroup, we used liverwort *SMXL* sequences to root the trees, consistent with the traditional view of land plant phylogeny (Qiu et al, 2006). More recent analyses have suggested hornworts might be the earliest diverging land plant lineage (Wickett et al, 2014), and we also able to root the tree with hornwort sequences without altering the topology of the tree. Consistent with both long-held and current notions of organismal phylogeny in land plants, the liverwort, moss, and hornwort *SMXL* clades are arranged as grade with respect to a large clade containing most tracheophyte sequences (Figure 6). As is common in such reconstructions, the large molecular rate heterogeneity amongst the lycophytes resulted in sequences from the Selaginellales grouping separately from the Isoetales and Lycopodiales, and with the latter two groups being drawn to the base of the tree (Cox et al, 2014). Given the known problems in this area, and the distribution of sequences, the most parsimonious explanation is lycophyte *SMXLs* form a monophyletic clade. Sequences from seed plants grouped into two super-clades which we denote as *SMAX1* and *SMXL4*. In our reconstruction, the Selaginellales *SMXLG/H* clades group with seed plant *SMXL4* and monilophyte *SMXLJ* with seed plant *SMAX1* (Figure 6). A strict reading of this phylogeny would imply a duplication at the base of vascular plants, followed by loss of one clade each in lycophytes and monilophytes. However, the most parsimonious explanation is that there was a duplication at the base of seed plants, and that the single lycophyte and monilophyte clades form a grade with respect to a monophyletic seed plant clade (Figure 7). We believe the data are most consistent with there being a complement of 1 *SMXL* gene in the last common ancestor of land plants, and with this basic complement being maintained during much of land plant evolution (Figure 7).

**Figure 6:**
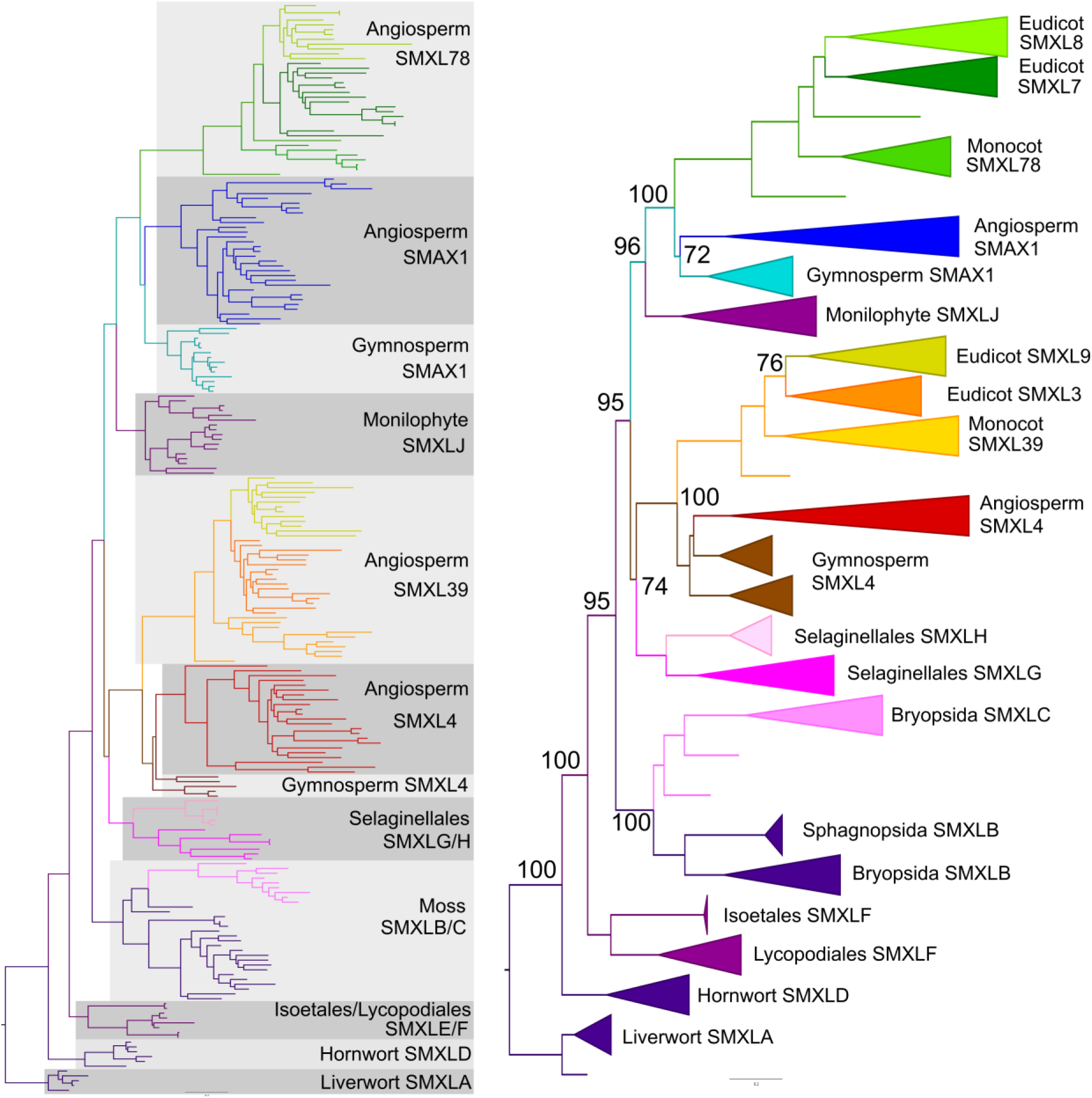
Phylogenetic analysis of the SMXL family. Nucleotide-level phylogenetic analysis implemented in PhyML on the *SMXL* family (226 sequences, 1431 characters). Trees were rooted with liverwort *SMXL* sequences. A) Phylogram showing the ‘most likely’ tree from PhyML analysis, labelled to show the high-order relationships between the major clades. B) Cladogram showing more detailed relationships between the clades listed in Table 1, with bootstrap values at key nodes.

**Figure 7:**
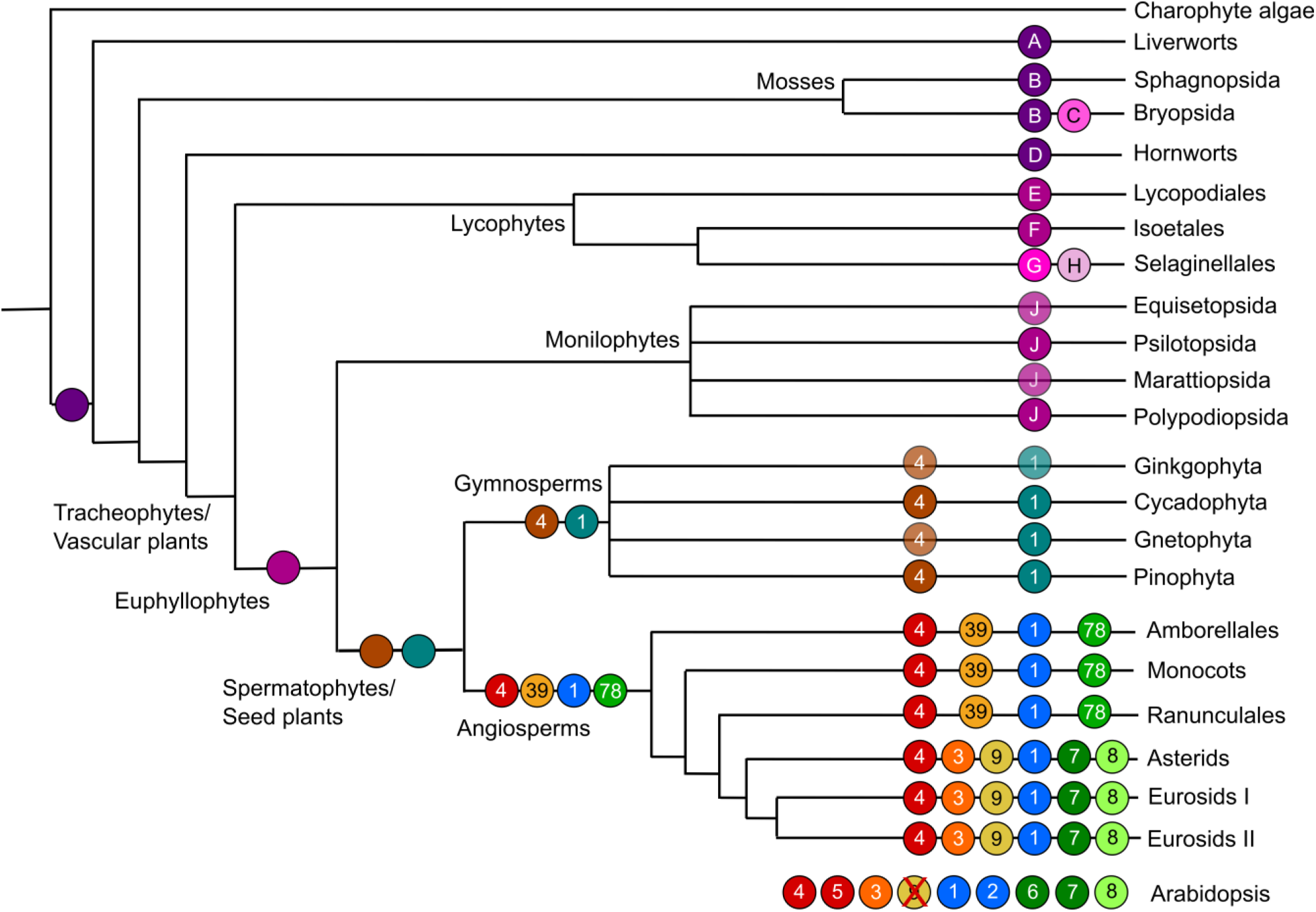
Reconstruction of SMXL evolution. Schematic depicting the complement of SMXL proteins in major land groups, and their inferred evolutionary origin. Each branch indicates a major land plant group; lycophytes, monilophytes and gymnosperms are further sub-divided into relevant orders/families/etc. The circles on each branch indicate the core complement of proteins in that group or sub-group. Clades which are inferred by parsimony are denoted with an opaque circle, and clade believed to have been lost are shown with a red cross. Letters and numbers in the circles indicate clade names. Circles without symbols at internal branching points represent the minimum inferred SMXL protein complement in the last common ancestor of each major land plant group.

In gymnosperms, there is one clade in each of the *SMAX1* and *SMXL4* super-clades, which we denote as *SMAX1* and *SMXL4* (Table 1). Our analysis suggests that there was a duplication in the *SMAX1* and *SMXL4* super-clades at the base of angiosperms, such that all angiosperms have at least four SMXL genes; *SMAX1* and *SMXL78* (*SMAX1* super-clade) and *SMXL39* and *SMXL4* (*SMXL4* super-clade) (Figure 7). Further duplications at the base of the eudicots has given rise to further sub-clades, *SMXL7* and *SMXL8* (within the *SMXL78* clade) and *SMXL3* and *SMXL9* (within the *SMXL39* clade), such that the basal complement of eudicots is 6 SMXL proteins (Figure 7). As previously described, Arabidopsis has 8 *SMXL* genes (Stanga et al, 2013); *SMXL3*, *SMXL8* and the pairs *SMAX1-SMXL2*, *SMXL6-SMXL7* and *SMXL4-SMXL5* which all represent recent duplications in the Brassicaceae; *SMXL9* has been lost from the lineage (Figure 7).

### SMXL proteins have a modular structure

In order to understand the possible functions of this extended set of SMXL proteins in seed plants, we looked more closely at the protein structure within the family (Figure 8; Supplemental data 10). As previously described at the N-terminus of each protein is a completely conserved double Clp domain (domain A; positions 1-164 in Arabidopsis SMAX1)(Stanga et al, 2013). There is almost no length variation at the N-terminus of the protein; only 4/226 proteins had any N-terminal extension. The double Clp domain consists of 5 sub-domains separated by less-conserved spacers of variable length. At the end of this domain is a clade-specific region (domain B; 165-213 in AtSMAX1), which aligns well within clades, but only very slightly across clades. Despite its variability, this region is always present. This is followed by a highly conserved region (domain C; 214-384 in AtSMAX1) of unknown function, which is present in all proteins except for SMXLB from mosses. Domain D (385-484 in AtSMAX1) is not as highly conserved but it does contain three highly conserved motifs (D1: CCxE/DC, D2: PSWLQ, D3: KLxELQKKWNxTCxSSLH). The domain is of similar size and structure in all proteins, and the motifs are in the same relative position across the whole family. After this domain there is another variable clade-specific region (Domain E: 485-604 in AtSMAX1) which aligns reasonably well within clades, but very poorly across the family as a whole. Despite this, domain E is present in all SMXL proteins, and contains one conserved motif (E1: SxVxTDLALGRS).

**Figure 8:**
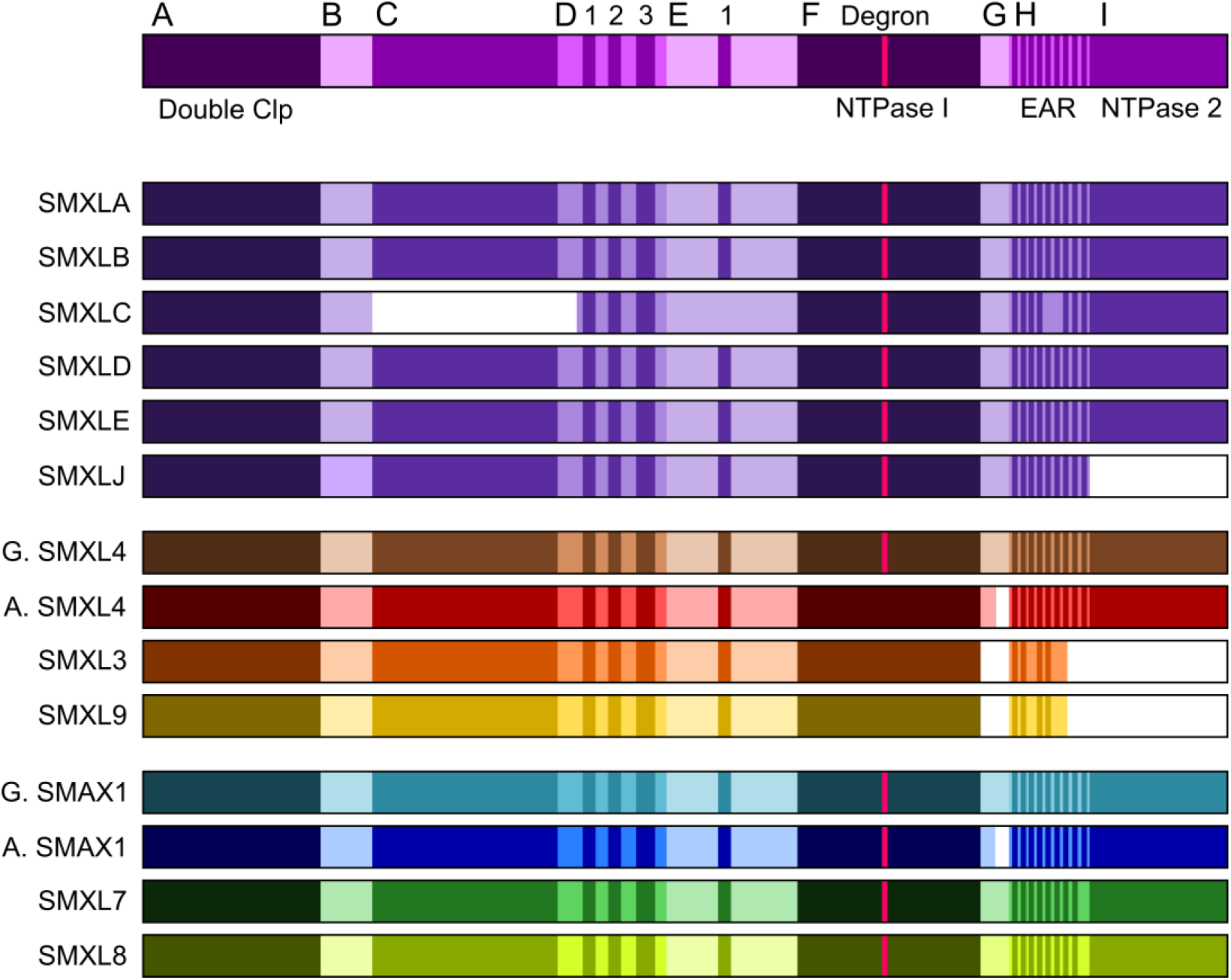
SMXL proteins have a modular structure. Top line: structure of a generic SMXL protein, showing the domains A-I, and their approximate level of sequence conservation between clades. Darker colours indicate regions that are highly alignable between clades, lighter colours indicate regions only alignable within clades. Conserved motifs within domains D, E and H are indicated. The magenta line within domain F indicates the FRGKT degron motif. Previously described features are indicated. Other lines: structure of different SMXL protein types, illustrating which features are conserved or absent. Proteins are coloured to match Figure 7.

Immediately after domain E, there is the previously described first NTPase domain (domain F; 605-773 in AtSMAX1), which is completely conserved, and which contains the FRGKT ‘degron’ motif needed for SL-induced degradation of SMXL7/D53 proteins. This is followed by a short clade-specific region (domain G: 774-800 in AtSMAX1) which is a similar size and structure in non-seed plants SMXLs, but is shorter or absent in seed plant SMXLs and does not align well between clades. Domain H (801-874 in AtSMAX1) consists of series of approximately 9 short alignable motifs, which occur in a conserved order, but with somewhat variable spacing. A few of these motifs are completely conserved, including the previously described EAR motif (H4), but some are missing in different clades. Finally, there is a second NTPase domain and a short C-terminal tail (domain I; 875-990 in AtSMAX1). Intriguingly, this domain is not present in monilophyte SMXLs, nor in angiosperm SMXL3 or SMXL9 proteins (Figure 8).

### Sub- and Neo-functionalization of SMXL proteins

SMXL3 and SMXL9 proteins thus show major innovation in protein structure relative to other SMXL proteins, lacking domains G and I, along with motifs H3 and H6-9. These data strongly suggest that they are neo-functional with regard to the ancestral SMXL function. Conversely, SMAX1, SMXL4, SMXL7 and SMXL8 proteins have broadly similar domain structure to SMXLs from non-seed plants. This raises interesting questions regarding which, if any of these proteins maintains the ancestral function of SMXL proteins, and whether we can therefore infer anything about this ancestral function.

To assess whether these proteins are likely to be neo- or sub-functionalized, we calculated pairwise identity scores between complete or near complete sequences in our alignment (167 sequences), for all the positions included in our phylogenetic analysis (477 amino acids). For every seed plant sequence, we calculated % identity with each SMXL protein from liverworts, mosses, hornworts, lycophytes and monilophytes (excluding the highly divergent SMXLB, SMXLG and SMXLH proteins). For each seed plant sequence we then calculated the average % identity with non-seed plant SMXLs. Finally, we then averaged these averages across each seed plant SMXL clade or sub-clade (Table 2; Supplemental data 11). Interestingly, this analysis showed that gymnosperm SMAX1 and SMXL4 proteins are equally identical to ancestral SMXL proteins (Table 2), suggesting that they may simply be sub-functionalized with respect to the ancestral function. Conversely, angiosperm proteins showed more differentiation from non-seed plant SMXLs, suggestive of neo-functionalization within angiosperm proteins (Table 2). Angiosperm SMAX1 proteins are the least differentiated from non-seed plant SMXLs (44.6% average identity). It could not be said that SMAX1 unambiguously preserves the structure of ancestral SMXLs, but compared to other angiosperms SMXLs, SMAX1 is the most likely candidate to maintain ancestral SMXL function.

**Table 2:** **SMXL protein identity comparison** Protein identity comparisons between different SMXL groups. Pairwise identity scores were calculated for each of 167 full-length/near full-length proteins in our alignment, across the 477 amino acids used for phylogenetic reconstruction. For each seed plant sequence, we then averaged their % identity across non-seed plant sequences from SMXLA, SMXLB, SMXLD, SMXLE, SMXLF and SMXLJ clades. We then averaged these scores to reach an average % identity per clade (2^nd^ column). Additionally, for each angiosperm SMXL4/SMXL39/SMXL3/SMXL9 sequence, we calculated % identity with gymnosperm SMXL4 sequences, then averaged across the clade (3^rd^ column). Finally, for each angiosperm SMAX1/SMXL78/SMXL7/SMXL8 sequence, we calculated % identity with gymnosperm SMAX1 sequences, then averaged across the clade (4th column).

|  | SMXLA-J | G. SMXL4 | G. SMAX1 |
| --- | --- | --- | --- |
| G.SMXL4 | 49.9 |  |  |
| A.SMXL4 | 40.0 | 50.9 |  |
| SMXL39 | 39.9 | 48.2 |  |
| SMXL3 | 41.6 | 50.7 |  |
| SMXL9 | 38.6 | 47.1 |  |
| G.SMAX1 | 49.1 |  |  |
| A.SMAX1 | 44.6 |  | 60.4 |
| SMXL78 | 34.3 |  | 41.0 |
| SMXL7 | 34.7 |  | 41.2 |
| SMXL8 | 34.1 |  | 40.9 |

### Specific strigolactone targets are an angiosperm-specific innovation

The SMXL78 clade in angiosperms consists of proteins that are degraded in response to strigolactones, and regulate a suite of developmental traits that are associated with strigolactones. These proteins are highly differentiated from non-seed plant SMXLs (Table 2), suggesting that they are likely to be neo-functional with regard to ancestral function. It is also notable that there are no separate SMXL78-like proteins in gymnosperms, suggesting that specific strigolactone targets represent a very recent evolutionary innovation. However, an alternative possibility is that gymnosperm SMAX1 proteins perform the role of both SMAX1 and SMXL7, and sub-functionalized into eu-SMAX1 and SMXL7 proteins in angiosperms. To examine this idea, we calculated the average identity for each angiosperm SMXL78 or SMAX1 protein with the SMAX1 proteins from gymnosperms, then averaged across each clade (Table 2; Supplemental data 11). SMXL78 proteins share about 41% identity with gymnosperm SMAX1, compared to 60% identity in angiosperm SMAX1 proteins. Taken together, the data strongly suggest that angiosperm and gymnosperm SMAX1 proteins have a conserved function, and that angiosperm SMXL7 proteins are neo-functional.

### Non-labile SMXLs are an angiosperm-specific innovation

In Arabidopsis, SMXL3, SMXL4 and SMXL5 have been characterized as non-labile, on the basis that they lack the FRGKT ‘degron’ motif that confers lability on SMXL7, and their increased stability relative to SMAX1 and SMXL7 (Wallner et al, 2017). This motif is broadly present in all non-seed plant SMXLs, and in SMAX1 and SMXL7/SMX8 proteins, but is absent from all SMXL3, SMXL4 or SMXL9 proteins in angiosperms (Figure 8; Supplemental data 11). Conversely, we observed that SMXL4 proteins from gymnosperms still retain the FRGKT motif. This suggest that non-lability is an angiosperm specific innovation. While SMXL4 proteins from gymnosperms are no more differentiated from ancestral SMXL proteins than gymnosperm SMAX1, angiosperm SMXL4 proteins are more highly differentiated proteins at the level of protein sequence (Figure 8, Table 2). Indeed, in terms of protein sequence in conserved regions, angiosperms SMXL4 is no more similar to gymnosperm SMXL4 than SMXL3/SMXL9 are (Table 2). Thus, although angiosperm SMXL4 does not have radical structural innovations, it may not have comparable functions to gymnosperm SMXL4 proteins. Taken together, the data suggests that seed plant SMXL4 proteins were originally sub-functionalized relative to the ancestral SMXL protein, but that the angiosperm members of this clade are now all likely to be at least partly neo-functional.

## DISCUSSION

### Ancient enzymes, ancient signals

A large and growing number of SL-type molecules have now been identified from root exudates (Yoneyama et al, 2017). Synthesis of SLs has been reported from across the land plant clade, including in liverworts (Delaux et al, 2012), mosses (Proust et al, 2011), lycophytes, gymnosperms and angiosperms (Yoneyama et al, 2017). This apparent broad distribution has been paradoxical when set against the apparent absence of core SL synthesis enzymes in many model species; for instance *Marchantia polymorpha* lacks a *CCD8* and *MAX1* orthologue, while *Physcomitrella patens* also lacks *MAX1* (reviewed in Waters et al, 2017). This has led to the suggestion of non-canonical synthesis pathways for SLs (Waldie et al, 2014; Waters et al, 2017). However, there appears to be a rather more straightforward answer to this paradox. Our re-examination demonstrates that the complete set of core SL synthesis enzymes, and potentially the recently identified LBO oxidoreductase, are broadly present across land plants and the antecessors of these enzymes are present in charophyte algae (Figure 9). Clearly, as with *M. polymorpha* and *P. patens*, individual species have lost genes, but taken as a group, we found all the core enzymes present in liverworts, mosses, lycophytes, gymnosperms and angiosperms. We only identified *CCD8* in hornworts, but this may be due to relatively low density of transcriptome assemblies in this group, especially if the tissues expressing SL synthesis genes were not sampled. In general, we struggled to identify *D27* and *CCD7* sequences from transcriptome assemblies, suggesting these genes have very low or very spatially restricted expression across land plant families.

**Figure 9:**
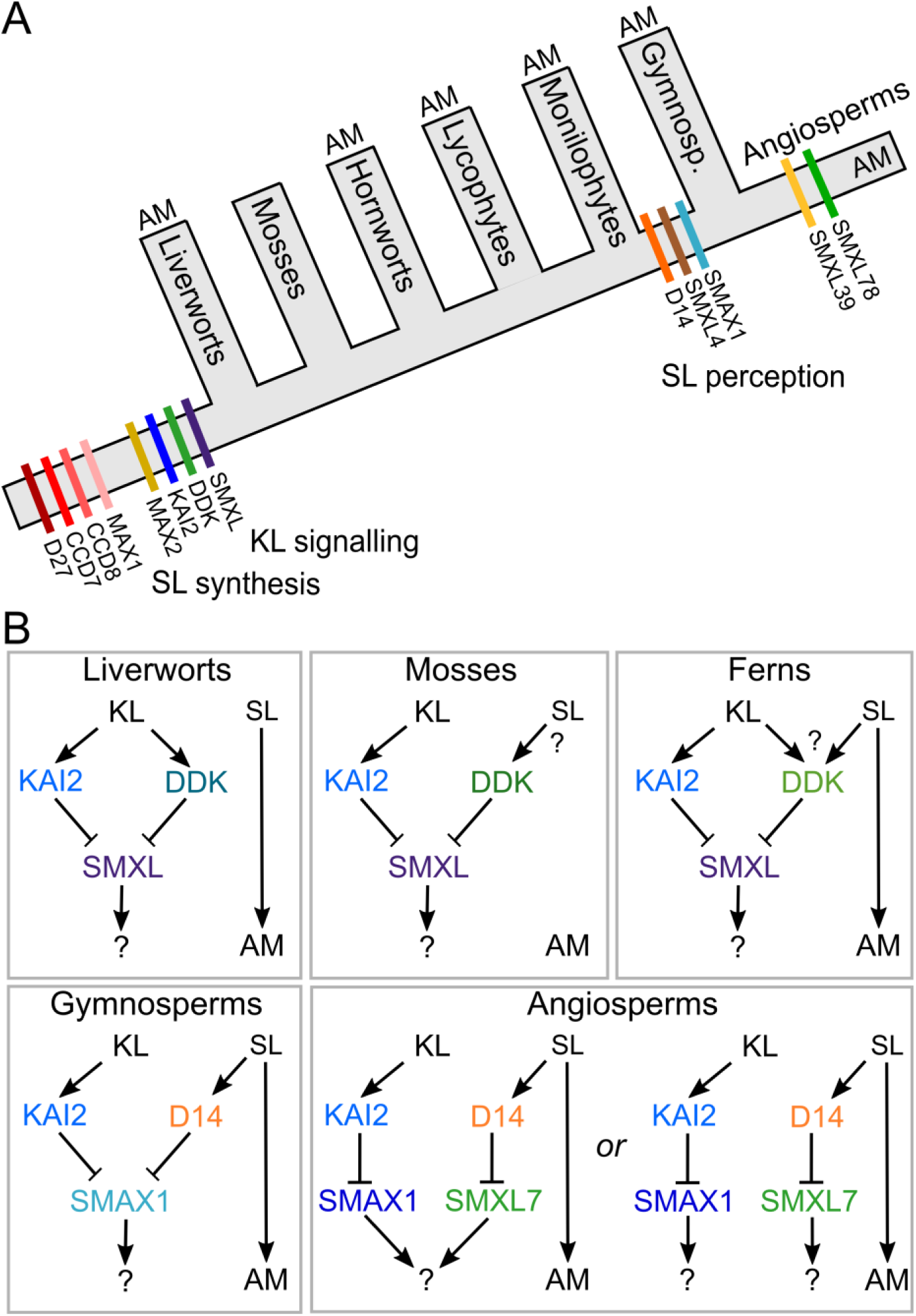
Models of SL evolution. A) Schematic of land plant evolution, showing key innovations in SL/KL synthesis and signaling. The organismal phylogeny depicted in grey follows traditional notions of land plant phylogeny (see Qui et al, 2006). ‘AM’ indicates that organisms in the group usually form mycorrhizal associations. B) Possible models of SL function throughout land plant evolution. In liverworts, there does not seem to be any canonical SL signaling. In mosses, SL perception may have evolved in the DDK lineage, and mycorrhizal associations have been lost. The status of SL and KL signaling in lycophytes and monilophytes is largely unclear. In gymnosperms, canonical SL signaling is present, but both SL and KL target the same SMXL protein for degradation. In angiosperms, SL and KL target separate SMXL proteins. It is unclear whether downstream targets are still shared, or have also separated.

Overall, the evolutionary distribution of SL synthesis enzymes is broadly consistent with the reported detection of SLs themselves. However, some caution is required, as it now appears that early mass spectrometry experiments have produced false positive signals for SLs in many species (reviewed in Yoneyama et al, 2017). Thus, while *P. patens* was previously found to produce several SLs, including some even in the absence of CCD8 (Proust et al, 2011), re-examination of this species failed to identify any SLs other than carlactone (Yoneyama et al, 2017). This is consistent with the presence of D27, CCD7 and CCD8 in *P. patens*, and the lack of MAX1. In a similar vein, *M. polymorpha* was previously reported to produce SLs (Delaux et al, 2012), but this seems unlikely given the lack of CCD8 and MAX1 in this species. Perhaps most pertinently, it was previously suggested that charophyte algae in the Charales produced the rather derived SL-type sorgolactone. Given the absence of true CCD7 and CCD8 enzymes in charophyte algae, and the failure to detect SLs in other charophytes, it seems plausible that the detection of sorgolactone in these species was a false positive. We suggest that SLs are only produced in land plants, consistent with the apparent evolution of true SL synthesis enzymes at the base of land plants (Figure 9).

### A late origin for strigolactone signalling?

Previous work showed that the evolution of eu-D14 type SL receptors occurred relatively recently in the seed plants (Bythell-Douglas et al, 2017); consistent with this, we find that the evolution of specific SL signalling targets is even more recent, occurring within the angiosperms. While it remains possible that ‘DDK’ proteins from lycophytes and monilophytes act as SL receptors (Bythell-Douglas et al, 2017), there is currently little good evidence that either of these groups show developmental responses to SLs. In liverworts, DDK proteins seem unlikely to act as SL receptors (Bythell-Douglas et al, 2017), while evidence in mosses is mixed; some DDK proteins might feasibly act as SL receptors (Lopez-Obando et al, 2016). Developmental responses to SL treatment have been claimed in *Chara corallina*, in liverworts and in mosses (Proust et al, 2011; Delaux et al, 2012), but these assays all used *rac*-GR24, and thus the responses cannot confidently be attributed to SL-like molecules. Only in *P. patens*, where development is altered in the absence of CCD8 is there evidence that development is regulated by SLs (Proust et al, 2011). The lack of apparent SL signalling mechanisms in non-seed plants has led to non-canonical signalling mechanisms being proposed (Bennett & Leyser, 2014).

Taking a parsimonious view, a more simple explanation would be that developmental responses to SLs are not an ancestral characteristic of land plants, consistent with the absence of SL signalling factors from non-seed plants. While this is perhaps a controversial viewpoint, it is worth exploring the ramifications of this idea. If SLs are not developmental regulators in non-seed plants, then why are they (apparently) synthesized in these species? The most obvious answer would be that SLs evolved firstly as rhizosphere signalling molecules, and would have subsequently been recruited as ‘internal’ hormones in seed plants by evolution of SL signalling. Indeed, the idea that SLs are ancestral rhizosphere signals has previously been suggested (Bouwmeester et al, 2007). In this respect, it is worth considering the case of *M. polymorpha*, which has lost SL synthesis enzymes relative to its close relatives *M. paleacea* and *M. emarginata* (Supplemental data 4, Supplemental data 6), and has also lost the ability to form mycorrhizal symbioses relative to these species. While cause and effect has not been established here, it is nevertheless unlikely to be coincidental. Mosses as a group have lost the ability to form mycorrhizal associations, and it may thus be that they have independently evolved the ability to use SL in the regulation of development. The main role of SLs in *P. patens* seems to be in the regulation of colony morphology and ‘quorum sensing’, and this can be seen as an alternative use of SLs as rhizosphere signals, rather than a secondary use as hormones. Mosses may have convergently evolved SL receptors from DDK proteins, though the evidence for this is mixed (Lopez-Obando et al, 2016, Bythell-Douglas et al, 2017), or may have completely separate SL receptors. More work is required to test the developmental roles of SLs in non-seed plants, but currently available evidence suggests that a late origin for SL signalling and developmental responses is a plausible hypothesis.

### An ancient KAI2-SMXL module

By contrast with SL signalling, there is little uncertainty regarding the origin of KAI2 signalling in land plants, since KAI2 proteins are present across the clade and proto-KAI2 proteins are found in charophyte algae (Bythell-Douglas et al, 2017). Here we have shown that SMXL proteins are also found throughout land plants, and that in non-seed plants, only a single SMXL type is found. We also show that these non-seed plant SMXLs have the ‘degron’ motif found in D53/SMXL7, suggesting that they may be degraded in a SCF^MAX2^-dependent manner. Given the ubiquity of KAI2 proteins, it seems most likely that SMXL proteins in non-seed plants are degraded in response to KAI2 activation, and indeed, we have shown that, of SMXL proteins found in angiosperms, nonseed plant SMXL proteins most closely resemble SMAX1, the presumptive target of KAI2 signalling. Although evidence is only circumstantial at this point, we believe that KAI2-induced degradation of SMXL proteins is an ancestral signalling mechanism in land plants.

We previously proposed that canonical SL signalling arose from neo-functionalization of KAI2-family receptors in seed plants, a process which was certainly complete by the last common ancestor of extant seed plants (Bythell-Douglas et al, 2017). However, gymnosperms only have two classes of SMXL protein, neither of which resembles SMXL7/D53 proteins, and our phylogenetic analysis clearly shows *SMXL7* arising from within the *SMAX1* lineage in angiosperms Thus, while SL and KL signalling have different target proteins in angiosperms, our analysis suggests they target the same protein for degradation in gymnosperms, and thus presumably regulate essentially the same downstream developmental processes (Figure 9). This raises the question as to whether KL and SL signalling also essentially regulate the same downstream processes in angiosperms, or whether the SL signalling pathway in angiosperms has completely novel targets relative to KL signalling.

### Other SMXL signalling modules

It has previously been observed that SMXL3/SMXL4/SMXL5 from Arabidopsis lack the conserved degron motif and are not degraded in response to KL or SL (Wallner et al, 2017), and we show here that this is a conserved feature of the SMXL3 and SMXL4 clades in angiosperms, as well as identifying a third clade of ‘non-degradeable’ SMXLs (SMXL9). We have also previously identified well-conserved clades of angiosperm D14/KAI2 family proteins (DLK2 and DLK3) that lack conserved residues needed for interaction with MAX2. It is thus tempting to speculate that DLK2/DLK3 proteins may regulate the activity of SMXL3/SMXL4/SMXL9 proteins, in response to unknown ligands, in a non-proteolytic manner. In this context it is notable that the evolution of DLK3 and SMXL9 appear to track each other to some extent. Both arise after the divergence of monocots from other angiosperms, and both have been lost from the Brassicaceae. We thus suggest that DLK3 and SMXL9 may be partners, and that DLK2 is a partner for either SMXL4 or SMXL3. While these proposals are currently speculative, these enigmatic members of the D14/KAI2 and SMXL families are starting to be characterized (Vegh et al, 2017; Wallner et al, 2017), and it will very interesting to gain further insights into their function and interaction.

## MATERIALS & METHODS

### Bioinformatic retrieval of *D14/KAI2* and *MAX2* sequences

Members of the *D27*, *CCD7*, *CCD8*, *MAX1*, *LBO* and *SMXL* families were identified by BLAST searches against complete, annotated genomes from two major sources: Phytozome (www.phytozome.net), or the genome portals for individual species. BLAST searches were performed using the coding sequences from *Arabidopsis thaliana*. Preliminary trees were assembled and used to guide the iterative interrogation of transcriptome databases, particularly those generated by the 1KP project (https://www.bioinfodata.org/Blast4OneKP/home). All sequences are listed in Supplemental data 12. For transcriptome datasets, we BLASTed each major taxonomic group separately. For non-annotated sequences from transcriptome datasets, we searched translations across all 6 reading frames to identify ORFs, and the longest ORFs were extracted for alignment.

### Alignment

Alignments were initially performed in BioEdit (Hall, 1997) using ClustalW (Thompson et al, 1994). Full length sequences from completed genomes were used for the initial alignment, which was manually refined as necessary. We then added sequences from transcriptome databases, many of which are incomplete, but the alignment of full length sequences provided a scaffold to align these sequences correctly. The resultant alignments are provided in Supplemental data 1,3,5-8,10. Pairwise protein identities were calculated using BioEdit. with the ‘Protein identity matrix’ function.

### Phylogenetic analysis

For each alignment family we performed preliminary phylogenetic analyses to explore the topology of the tree and the effect of inclusion or exclusion of various groups of sequences. We trimmed all alignments to remove poorly conserved regions. Final maximum likelihood analyses were performed with PhyML (Guindon et al, 2010) using a GTR+G+I model on nucleotide level data. Trees were visualized and modified using FigTree 1.4.2.

## Authors’ contributions

TB conceived and designed the study. CW analysed sequence data and performed phylogenetic reconstructions. CW & TB wrote the manuscript.

## Acknowledgements

We gratefully acknowledge the use of sequence data generated by members of the 1000 Plants (1KP) initiative, and in particular Gane Ka-Shu Wong, Carl Rothfels, Sean Graham, Dennis Stevenson, Michael Melkonian, Barbara Surek, Jim Leebens-Mack, Michael Deyholos, Douglas Soltis and Pamela Soltis. TB is supported by RG160845 from the Royal Society and BB/R00398X/1 from BBSRC.

---

**Supplemental data 1: D27 alignment**

**Supplemental data 2: LBO sequence identity matrix**

**Supplemental data 3: CCD7 alignment**

**Supplemental data 4:**
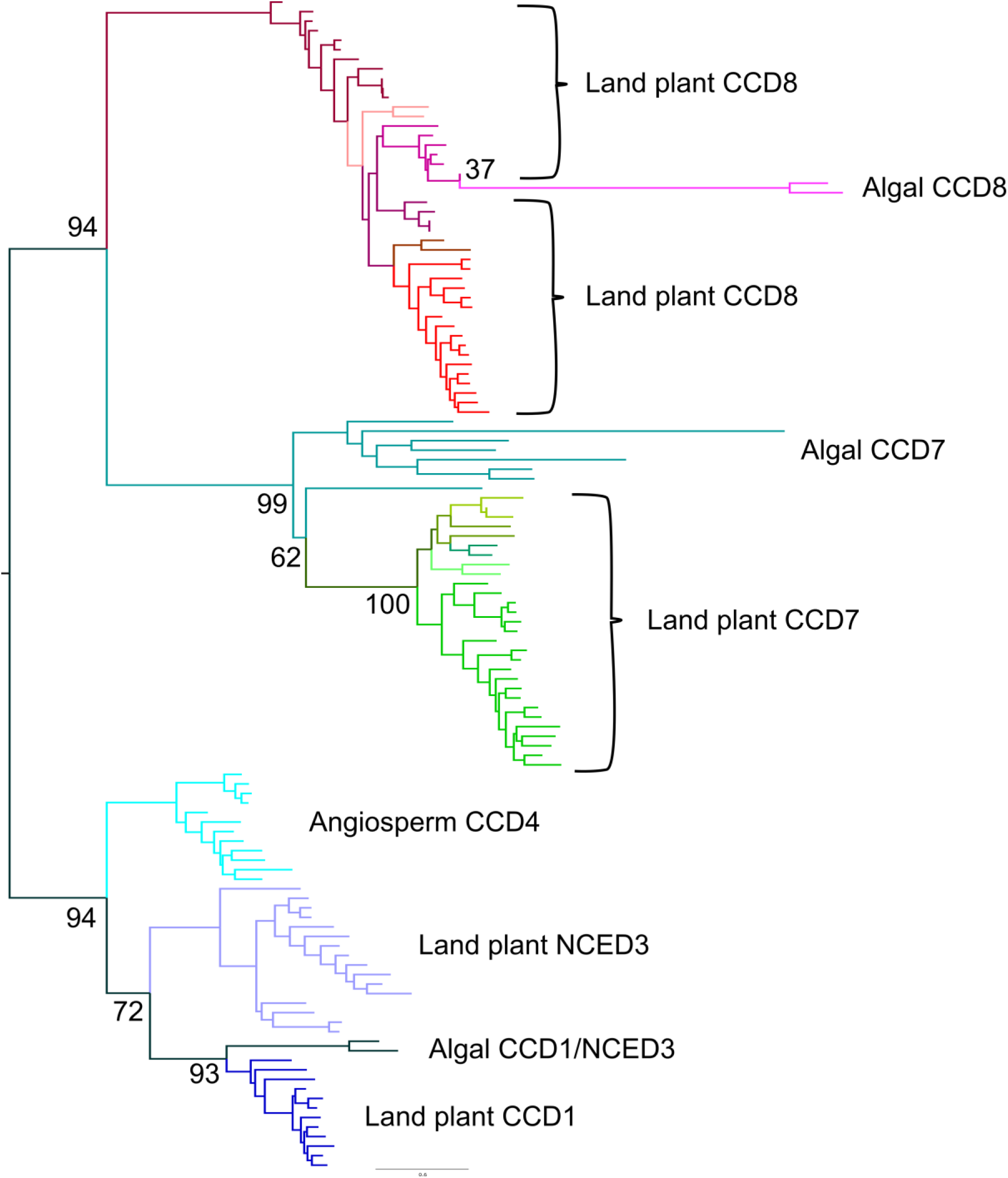
CCD phylogeny. Nucleotide-level phylogenetic analysis implemented in PhyML on the *CCD* family (123 sequences, 919 characters). The tree was rooted with an algal sequence. Phylogram showing the ‘most likely’ tree with bootstrap values at key nodes.

**Supplemental data 5: CCD alignment**

**Supplemental data 6: CCD8 alignment**

**Supplemental data 7: MAX1 alignment**

**Supplemental data 8: LBO alignment**

**Supplemental data 9: LBO sequence identity matrix**

**Supplemental data 10: SMXL alignment**

**Supplemental data 11: SMXL sequence identity matrix**

**Supplemental data 12: List of sequences**

